# Discrete action control for prosthetic digits

**DOI:** 10.1101/2020.03.25.007203

**Authors:** Agamemnon Krasoulis, Kianoush Nazarpour

**Affiliations:** School of Engineering, Newcastle University, Newcastle-upon-Tyne, NE1 7RU, UK; Biosciences Research Institute, Newcastle University, Newcastle-upon-Tyne, NE2 4HH, UK

## Abstract

**Objective:** We aim to develop a paradigm for simultaneous and independent control of multiple degrees of freedom (DOFs) for upper-limb prostheses.

**Approach:** We introduce *action control*, a novel method to operate prosthetic digits with surface electromyography (EMG) based on multi-label, multi-class classification. At each time step, the decoder classifies movement intent for each controllable DOF into one of three categories: open, close, or stall (i.e., no movement). We implemented a real-time myoelectric control system using this method and evaluated it by running experiments with one unilateral and two bilateral amputees. Participants controlled a six-DOF bar interface on a computer display, with each DOF corresponding to a motor function available in multi-articulated prostheses.

**Main results:** We show that action control can significantly and systematically outperform the state-of-the-art method of position control via multi-output regression in both task- and non-task-related measures. Improvements in median task performance ranged from 20.14% to 62.32% for individual participants. Analysis of a post-experimental survey revealed that all participants rated action higher than position control in a series of qualitative questions and expressed an overall preference for the former.

**Significance:** Action control has the potential to improve the dexterity of upper-limb prostheses. In comparison with regression-based systems, it only requires discrete instead of real-valued ground truth labels, typically collected with motion tracking systems. This feature makes the system both practical in a clinical setting and also suitable for bilateral amputation. This work is the first demonstration of myoelectric digit control in bilateral upper-limb amputees. Further investigation and pre-clinical evaluation are required to assess the translational potential of the method.

## 1. Introduction

Modern hand prostheses offer the potential of partially restoring the functionality of a missing upper-limb. They are typically controlled by muscular activity signals recorded on the skin surface using electromyogram (EMG) signals. Nowadays, there exist commercial systems allowing for grip selection and activation based on machine learning algorithms. Nevertheless, from a technical perspective the holy grail of upper-limb prosthetics research is the simultaneous and independent control of multiple degrees of freedom (DOFs), including wrist and digit artificial joints [1]. This currently seems as the only way to approximate the remarkable dexterity of the human hand, which is still considered as the nature’s most versatile end-effector [2].

Significant efforts have been made towards achieving simultaneous, multi-DOF myoelectric control. The term *proportional control* is often used to describe the feasibility of controlling one or more prosthesis output(s) in a continuous space [3]. To that end, many research groups, including us, have used multi-output regression-based algorithms to map features extracted from surface or intramuscular EMG channels onto wrist [4–8] and/or finger position [9–14], velocity [15, 16], and force trajectories [17–20]. For prosthetic digit control, several studies have shown premise in decoding position/velocity offline [9, 11–13, 15, 16], however, only a smaller number have achieved real-time digit control in amputees [10,14,21]. This indicates that, to date, reconstruction of position/velocity trajectories from surface EMG signals in real-time remains a challenging problem.

In this work, we introduce *action control*, a novel paradigm for simultaneous and independent control of multiple prosthetic digits. Our approach simplifies the movement intent decoding component and makes the system more practical. In the heart of the algorithm lies a multi-label, multi-class classifier, which decodes movement intent for each controllable DOF into one of three classes (i.e., *actions*): open, close, or stall (i.e., no movement). A schematic diagram of this concept is shown in figure 1. From a control theory point of view, action control can be viewed as an extreme, discretised case of velocity control. The relationship between digit positions, velocities and actions is illustrated in figure 2. Despite using a classifier, action control allows digit positions to take values within a large set that is defined by the action step (i.e., position increase/decrease resolution). That is, by selecting a relatively small step, digit movement essentially becomes continuous. One practical advantage of this method over regression-based position control is that it allows for collection of noise-free ground truth labels, without relying upon the use of motion tracking hardware (e.g., data gloves).

**Figure 1:**
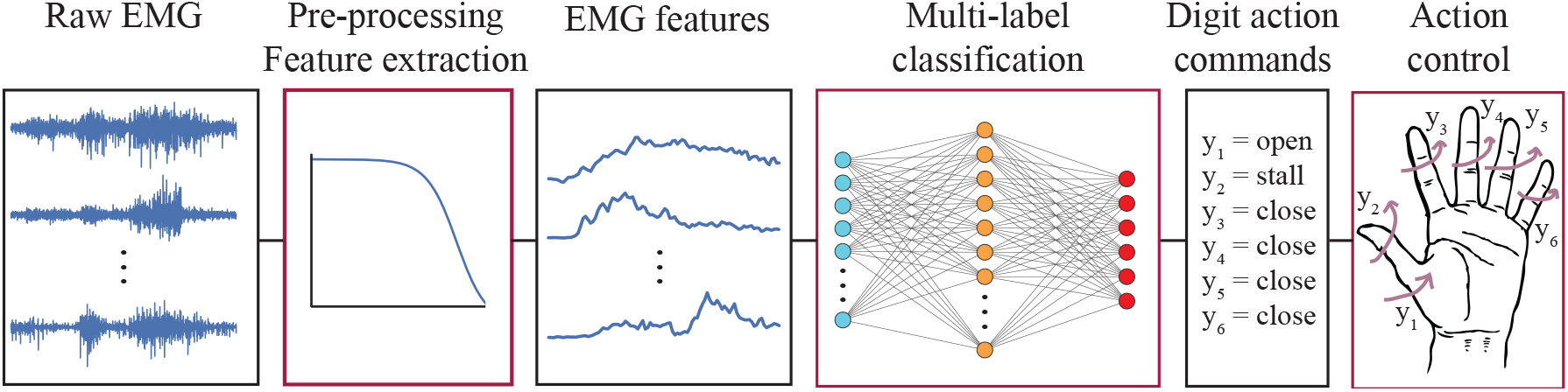
Action control concept. At each time step, raw EMG signals are pre-processed and fed as inputs into a multi-label classifier with six discrete outputs. These can take one of three possible values (i.e., *actions*): open, close or stall (i.e., no movement). Each output controls one DOF available in multi-articulated prosthesis: thumb rotation, and flexion/extension of the thumb, index, middle, ring, and little digits.

**Figure 2:**
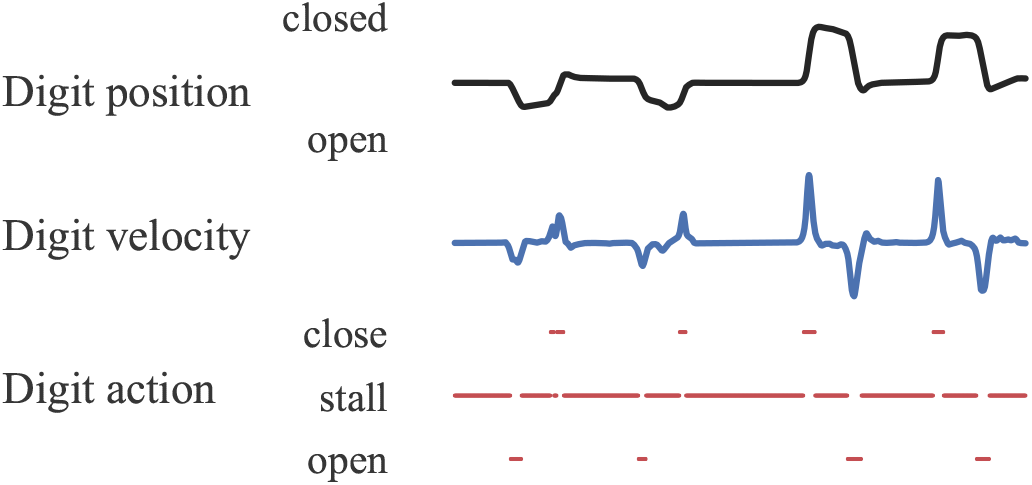
Relationship between digit position, velocity and action. Action control can be viewed as a discretised version of velocity control. Note that digit positions/velocities are not required for action control.

In a proof-of-concept study, we have previously demonstrated that users can employ the action control scheme to achieve comparable performance to joint angle (i.e. position) control in a robotic hand tele-operation task with an instrumented data glove [22]. In addition, we have performed a systematic offline investigation of multi-label classification for digit action decoding [23]. Here, we provide for the first time a real-time implementation of the algorithm for myoelectric digit control. We test the performance of our approach with one unilateral and two bilateral amputees and benchmark it against the state-of-the-art in myoelectric digit control, that is, regression-based joint position control. We discuss the advantages of our method over the regression-based approach and propose ideas for further improvement and pre-clinical evaluation. To our knowledge, this is the first demonstration of real-time, multi-DOF myoelectric digit control in bilateral upper-limb amputees.

## 2. Methods

### 2.1. Participant recruitment

Three transradial (i.e., below-elbow) amputee volunteer participants were recruited (two female, one male; median age 59). Participants 1 and 2 were bilateral amputees and participant 3 was unilateral. Paritcipant 1 performed the experiment using their right side. Participant 2 took part in two sessions on the same day using alternate sides (i.e., right one in the morning session and left one in the afternoon session). Depending on the side used in each session, the participant is referred to as 2R or 2L (right or left side, respectively). All participants had normal or corrected to normal vision and reported being able to discriminate red from blue (i.e., no colour blindness). Detailed demographic information about the participants is provided in Table 1.

**Table 1:** Participant demographic information.

Abbreviations: P, participant; G, gender; A, age; AT, amputation type; AS, amputated side; AC, amputation cause; YSA, years since amputation; FL, forearm length (cm); FC, forearm circumference (cm); Transr., transradial.
| P | G | A | AT | AS | AC | YSA | FL | FC |
| --- | --- | --- | --- | --- | --- | --- | --- | --- |
| 1 | F | 67 | Transr. | Bilateral | Sepsis | 5 | 13 | 22 |
| 2R | F | 42 | Transr. | Bilateral | Sepsis | 6.5 | 15 | 20.5 |
| 2L |  |  |  |  |  |  | 15 | 20.5 |
| 3 | M | 59 | Transr. | Right | Epithelioid sarcoma | 18 | 17 | 27 |

All experimental procedures were in accordance with the Declaration of Helsinki and approved by the local ethics committee at the Faculty of Science, Agriculture and Engineering (SAgE), Newcastle University (#17-NAZ-056). All participants read an information sheet and gave written informed consent prior to the experimental sessions.

### 2.2. Apparatus

The apparatus used in the study comprised a surface EMG recording system and a state-of-the-art, multi-articulated prosthetic hand.

#### 2.2.1. Surface EMG recording system

We recorded myoelectric signals using a Delsys^®^ Trigno™ wireless system. We used 16 EMG sensors, which included 12 standard Trigno and four Trigno Mini sensors. Prior to electrode placement, we cleansed participants’ skin using 70% isopropyl alcohol swabs. We then placed the EMG sensors in two rows of eight equidistant electrodes below the elbow without targeting specific muscles. In each row, we placed six standard and two mini sensors. We attached the transmitter units for the mini sensors on the participants’ skin above the elbow as shown in figure 3(a). Upon visual inspection of the signal quality of all EMG channels, we secured the electrode positions using adhesive elastic bandage. For data acquisition, we used proprietary software provided by the manufacturer. The sampling rate for all channels was fixed at 2 kHz.

**Figure 3:**
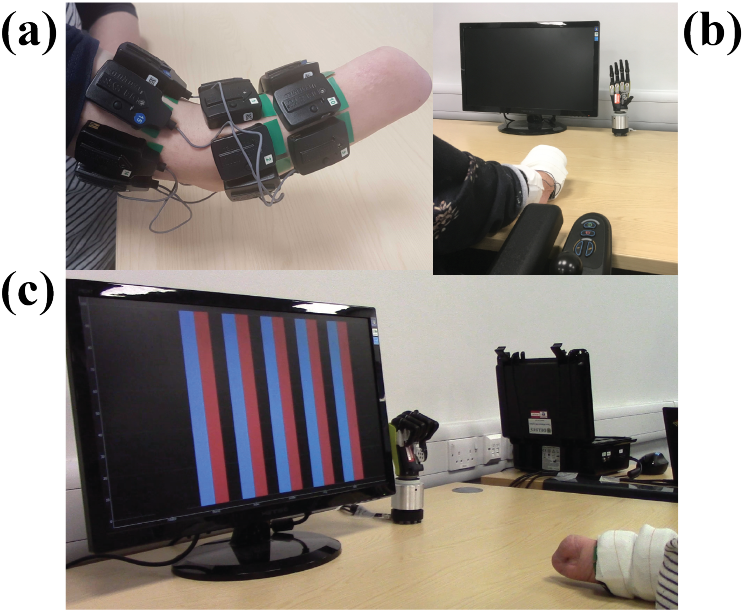
Experimental protocol. **(a)** Electrode placement for one participant. Sixteen surface EMG electrodes were placed below the elbow arranged in two rows of eight electrodes. A combination of classic Delsys Trigno and Trigno Mini electrodes were used. The transmitter components for the Mini sensors were placed above the elbow. **(b)** Training data collection. Participants performed imaginary movements of single-digit and full-hand grips that were instructed on the prosthesis. **(c)** Real-time decoding. Participants used their muscles to control the position of six bars on a screen. Target postures were instructed on the prosthesis prior to the start of the trial. The target posture for the shown trial was the “lateral” grip.

#### 2.2.2. Prosthetic hand

We used the multi-articulated Össur^®^ Robo-limb hand to demonstrate target postures to participants. This model, which is almost identical to the commercial i-Limb™ Ultra Revolution prosthesis, comprises six motors controlling the following DOFs: thumb rotation (i.e., opposition/reposition), and thumb, index, middle, ring and little finger flexion/extension. Throughout the experiments, the hand was placed on a desktop base and powered via an external power supply unit (7.4 V/7 A). The hand was controlled by a laptop computer using a CAN bus connection (baud rate: 1 MHz). We used a right-hand model for participants 1, 2R and 3 and a left-hand model for participant 2L.

### 2.3. Experimental paradigm

Following EMG sensor placement, participants sat comfortably on an office chair and rested their arm on a table. Experimental sessions comprised two phases: *training data collection* and *real-time myoelectric control*. The two sessions were interleaved with a 10-minute break, during which participants were allowed to stand up and/or move freely in the lab space.

#### 2.3.1. Training data collection

In the first part of the experiment, participants performed a series of imaginary finger movements that were instructed to them on the prosthesis as shown in figure 3(b). We included a combination of single-finger and full-hand grip patterns: thumb opposition/reposition; thumb, index, middle, ring and little finger flexion/extension; cylindrical and lateral grip opening/closing; and rest (i.e., no movement). We instructed participants to perform the exercises smoothly and follow the prosthesis movement as closely as possible.

For each exercise, the opening/closing, opposition/reposition or flexion/extension parts were separate. Thus the total number of exercises, including rest, was 17. Each exercise was repeated 12 times and each repetition lasted 1.3 s, which was the time required for the prosthesis to complete the demonstration at a moderate speed. Participants were given 2 s of rest in-between consecutive trials.

#### 2.3.2. Real-time myoelectric control

In the second part of the experiment, participants were instructed to use their muscles to control a six-dimensional bar interface as shown in figure 3(c). Visual feedback was continuously provided at an update rate of 15.625 Hz (section 2.4) on a 17” LCD flat panel display positioned 1 m in front of the participants.

An illustration of the experimental task is given in figure 4. Each pair of bars corresponded to one of the following DOFs: thumb opposition/reposition and thumb, index, middle, ring, and little finger flexion/extension. The blue bars were controlled by the participants and showed *actual* positions of the six DOFs. The red bars were kept fixed within the trial and corresponded to *target* positions. Participants were instructed to match the blue bars to the red ones as closely as possible.

**Figure 4:**
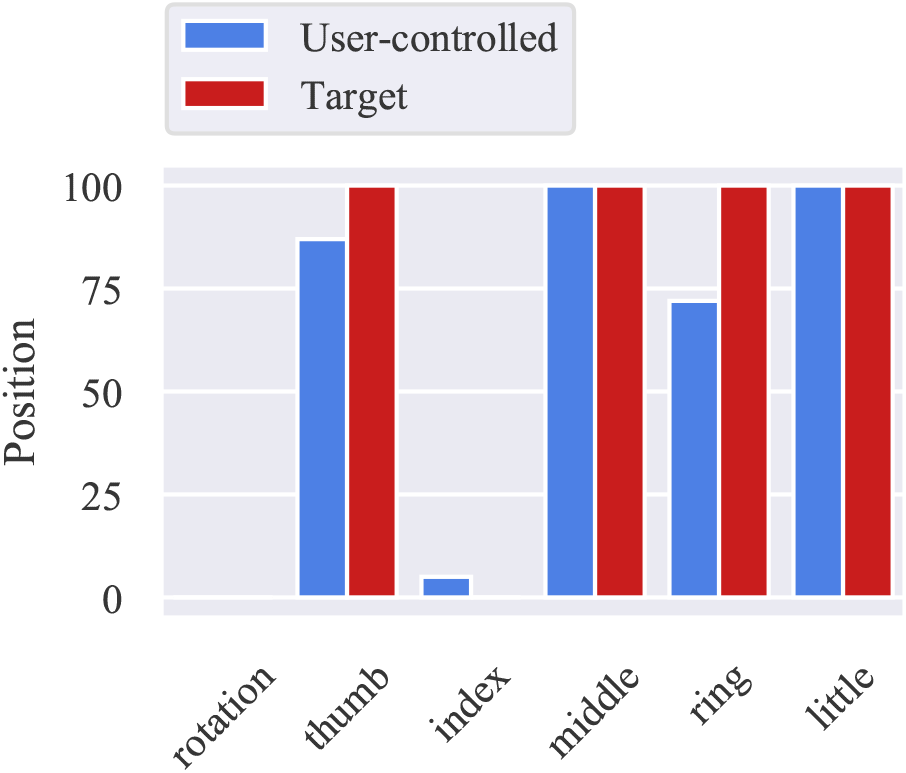
Real-time control task. Participants had to use their muscles to control the position of six bars on the screen, each corresponding to one DOF (from left to right: thumb rotation, thumb, index, middle, ring, and little finger flexion/extension). For each DOF, blue and red bars indicate the user-controlled and target positions, respectively. The shown example corresponds to the index pointer with the following target positions: thumb, middle, ring, and little digits fully flexed (i.e., target bars 2, 4, 5 and 6 in maximum position), index finger fully extended, and the thumb rotated fully reposed (i.e., target bars 1 and 3 in minimum position).

The following target movements were included: full thumb reposition; full flexion of the thumb, index, middle, ring, and little fingers; cylindrical, lateral, and tripod grips; and index pointer. Participants performed 10 blocks of trials for each of two control conditions (section 2.5), that is, a total of 20 blocks. Every target posture was included exactly once within a single block in a pseudo-randomised order.

Prior to the start of a trial, the target posture was demonstrated on the prosthesis. Once the demonstration was complete, an audio cue (sine wave; frequency 1000 Hz; duration 200 ms) initiated the start of the *preparation phase* of the trial and the bar interface appeared on the display (figure 4). Participants were then given 5 s to match the controlled posture (i.e., blue bars) to the target (i.e., red bars) as closely as possible. At the end of this period, a second cue indicated the start of the *evaluation phase* of the trial, which lasted for 1 s. During the evaluation phase, participants were instructed to hold the performed posture. At the end of the trial, the bar interface disappeared from the display and participants received a score characterising their performance during the evaluation phase (section 2.7). The prosthesis was also reset to the neutral (i.e., resting) posture. Consecutive trials were interleaved with 2.5 s of rest.

### 2.4. Signal pre-processing

Data were acquired and processed every 64 ms, which translates into 128 new samples/channel at a 2 kHz sampling rate. The data processing and display update rate thus was 1/0.064 s = 15.625 Hz. We processed EMG signals using a 128 ms sliding window (i.e., 50% overlap). We extracted two EMG features from each channel, namely, waveform length and log-variance [5]. We based our selection on previous findings showing that these features are effective both for multi-output regression and multi-label classification [14, 23]. We used the same EMG processing pipeline for the two control schemes presented in the following section.

### 2.5. Control schemes and decoder training

We implemented the *action control* method and compared it with the state-of-the-art approach of *position control*. The two algorithms are based on multi-label classification and multi-output regression, respectively.

All models were trained during the interval between the data collection and real-time control parts using subject-specific data. We discarded the first two repetitions for each exercise to avoid introducing noise in the training set, whilst participants familiarised themselves with a new exercise. Thus, 10 repetitions for each exercise were included in the training set for each participant.

#### 2.5.1. Action control

With action control, participants controlled the bar heights on the display by taking at each time step discrete actions in order to reach the desired positions. With this control mode, the EMG features were mapped onto a discrete six-dimensional vector using multi-label classification as shown in figure 1. Each element in the output vector corresponded to one of the six available DOFs and could take one of three values: open, close or stall (i.e., no movement). We set the action step for the “open” and “close” commands such that a single-DOF movement from the bottom to top position, or vice-versa, would require 1.5 s. This translated in using an action step of 0.043. Based on previous findings [23], we trained six independent linear discriminant analysis (LDA) classifiers, one for each available DOF.

We post-processed predictions using a confidence rejection strategy based on false positive rate minimisation [24]. With this method, class-specific thresholds were identified for each output using training data and 10-fold cross-validation, such that the false positive rate did not exceed a global cutoff threshold (set a priori to *θ*_*c*_ = 0.2). During inference (i.e., real-time control), predictions that were made with posterior probabilities smaller than the class-specific thresholds were discarded by the controller. Let us illustrate this with an example. Assume that the for the *j*^*th*^ DOF the rejection threshold for the “open” class is 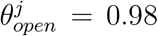. Also assume that at time step *t*, this DOF is stalled. If at step *t* + 1 the prediction for the same DOF is “open”, and the associated posterior probability is *p* = 0.9, then the controller will discard the new prediction and the DOF will hold its previous state, that is, it will remain stalled. For a more detailed description of this confidence-based rejection strategy, we refer the reader to our previous work [24].

#### 2.5.2. Position control

With position control, participants directly controlled the bar heights on the display. That is, the six-dimensional position trajectories were reconstructed from EMG features using multi-output regression and were normalised between 0 (i.e., digit fully extended or thumb fully reposed) and 1 (i.e., digit fully flexed or thumb fully opposed). We used standard linear regression models to map EMG features onto bar positions and post-processed predictions using single exponential smoothing [14]. We set the smoothing parameter a priori to *a* = 0.05.

### 2.6. Training data labelling

To train myoelectric decoders, ground truth data are required for both regression and classification tasks. In the former case, ground truth labels are real-valued, whereas in the latter they are discrete. In our experiments, labels were acquired using the prosthesis demonstrations presented to participants during the data collection phase.

For training regression models (i.e., position control), for which real-valued ground truth labels are required, we adopted the following strategy: at the start and end of each trial, each of the six DOFs are in one of the two extreme positions (i.e., thumb rotator fully opposed/reposed and each digit fully flexed/extended). Hence, the ground truth starting and ending points are known. Within the trial, the positions of those DOFs that are involved in the specific exercise are linearly interpolated between the respective starting and ending points. On the other hand, ground truth positions for those DOFs that are not involved in the trial movement are kept fixed and equal to the starting positions. To illustrate this approach, consider as an example the thumb opposition exercise. Assume ***y*** ∈ ℝ^6^ denotes the position vector with the first element corresponding to thumb rotation, the second element corresponding to thumb flexion and so on. At the start of the trial, all digits are fully extended and the thumb rotator is fully reposed. Thus, the ground truth starting point is ***y*** = [0, 0, 0, 0, 0, 0]^T^. At the end of the trial, only the thumb rotator has moved to the fully opposed position. That is, the ground truth ending point is ***y*** = [1, 0, 0, 0, 0, 0]^T^. The position of the thumb rotator is linearly interpolated within the trial, which lasts for 1.3 s. Note that this is exactly the time it takes for the prosthesis to demonstrate the exercise. For instance, at *t* = 0.65 s, the ground truth position is approximated by ***y*** = [0.5, 0, 0, 0, 0, 0]^T^.

Data labelling for multi-label classification models (i.e., action control) is simpler, since in this case ground truth labels are discrete. That is, 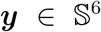, where 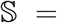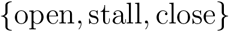 and we assume the same ordering in ***y*** as in the regression example above. For classification, however, we only need to know which DOFs are moving within a trial and in which direction. Considering again the thumb opposition example as above, the ground truth vector for the whole duration of trial is ***y*** = [close, stall, stall, stall, stall, stall]^T^.

### 2.7. Performance evaluation

At the end of each control trial, participants received a score on the screen characterising their performance during the evaluation phase of the trial. The score ranged from 0% to 100% and was based on the multivariate median absolute error *MAE_mv_* between the target and performed postures. This is defined as follows:

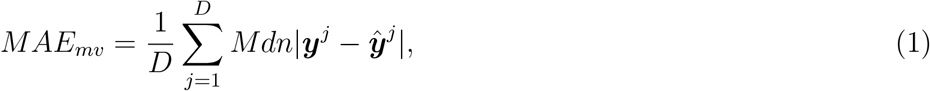

where ***y***^*j*^ ∈ ℝ^*N*^ and 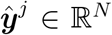 denote the vectors containing the target and performed position trajectories during the evaluation phase for the *j*^th^ DOF, *N* = 16 is the number of display updates (i.e., time steps) in the evaluation phase, *D* = 6 is the number of controllable DOFs, and *Mdn* denotes the median operator. The target position trajectories ***y***^*j*^ are known and constant (i.e., equal to the heights of the red bars in figure 4). On the other hand, the controlled position trajectories 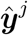 (i.e., the heights of blue bars) are variable, as they are controlled by the participant.

We transformed *MAE*_*mv*_ into an intuitive for the user performance score with the following principles in mind: 1) the score range should be from 0% to 100%; 2) a perfect match between the target and performed postures should result in a 100% score; and 3) a very poor or random prediction, defined as *MAE*_*mv*_ > 0.5, should yield a 0% score. Taking the above into consideration, we implemented this transformation as follows:

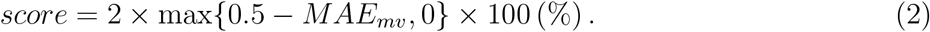

To assess offline decoding performance on the training set, we used 10-fold cross-validation. Within each iteration, nine out of 10 repetitions from all exercises were used for training and the remaining repetition was used for testing. We report the average performance across folds. For multi-output regression we use the multivariate coefficient of determination (R^2^) score and for multi-label classification we report macro-average— both across outputs and class labels— F1-score (i.e., harmonic mean of precision and recall).

### 2.8. Non-tasked-related measures

In addition to control performance, we assessed the two control schemes in terms of two non-task-related measures: *muscle contraction intensity* and *output stability*. For the former, we computed the raw EMG power across all channels during whole trial duration. For the latter, we estimated average output variability (i.e., standard deviation) during the trial evaluation phase.

### 2.9. Post-experimental questionnaire

At the end of each experimental session, participants were asked to respond to a short questionnaire. Concretely, they were asked to rate the two control schemes (i.e., action and position control) on a scale from 1 (strongly disagree) to 5 (strongly agree), based on the following statements:

i. the control interface was easy;
ii. the control interface was intuitive;
iii. I found it easy to adapt to the control interface.

Half scores, for example 3.5, were also allowed. Participants were finally asked to indicate which control interface they preferred overall. Participant 2 responded to the questionnaire twice, once after each experimental session, and the respective scores were averaged.

### 2.10. Implementation

The experiment was implemented in `Python 3` and all sessions were run on a standard laptop computer (16 GB, i7 @ 2.70 GHz). Data collection, processing, and the real-time myoelectric control interface were implemented using the `axopy` library [25] and custom-written code. Model training and inference were performed using the `scikit-learn` library [26].

### 2.11. Statistical analysis

We used two-sided Wilcoxon (i.e., non-parametric) tests throughout our analysis to perform comparisons between action and position control. We performed statistical comparisons at the participant level and used the Holm-Bonferroni correction method to account for multiple comparisons. The target presentation order was identical for the two conditions and, hence, all measurements were paired. The condition order was counter-balanced: participants 1 and 2L first performed the task with action and then position control, and participants 2R and 3 with the reversed order. We performed all statistical analyses in Python using the Pingouin library [27].

## 3. Results

### 3.1. Offline training and analysis

The results of the offline analysis performed as part of model training are presented in Table 2. The median macro-average F1-score for multi-label classification (i.e., action control) was 0.80 (ranging from 0.74 to 0.82) and the median R^2^ score for multi-output regression (i.e., position control) was 0.46 (ranging 0.38 to 0.51).

**Table 2:** Offline performance

Abbreviations: P, participant; $R^2$ , coefficient of determination (multi-output regression); F1, macro-average F1-score (multi-label classification)
| P | $R^2$ | F1 |
| --- | --- | --- |
| 1 | 0.51 | 0.82 |
| 2R | 0.38 | 0.74 |
| 2L | 0.40 | 0.78 |
| 3 | 0.51 | 0.82 |

### 3.2. Real-time control performance benchmarks

Performance results for the real-time control experiment are presented in figure 5. For each participant, performance scores from all trials are aggregated and summarised using box and violin plots. All participants achieved significantly higher performance with action than position control (P1, *MD* = 20.14, *p* < 10^−2^; P2R, *MD* = 52.63, *p* < 10^−13^; P2L, *MD* = 47.23, *p* < 10^−10^; P3, *MD* = 62.32, *p* < 10^−13^; *MD* denotes median difference between the two control schemes).

**Figure 5:**
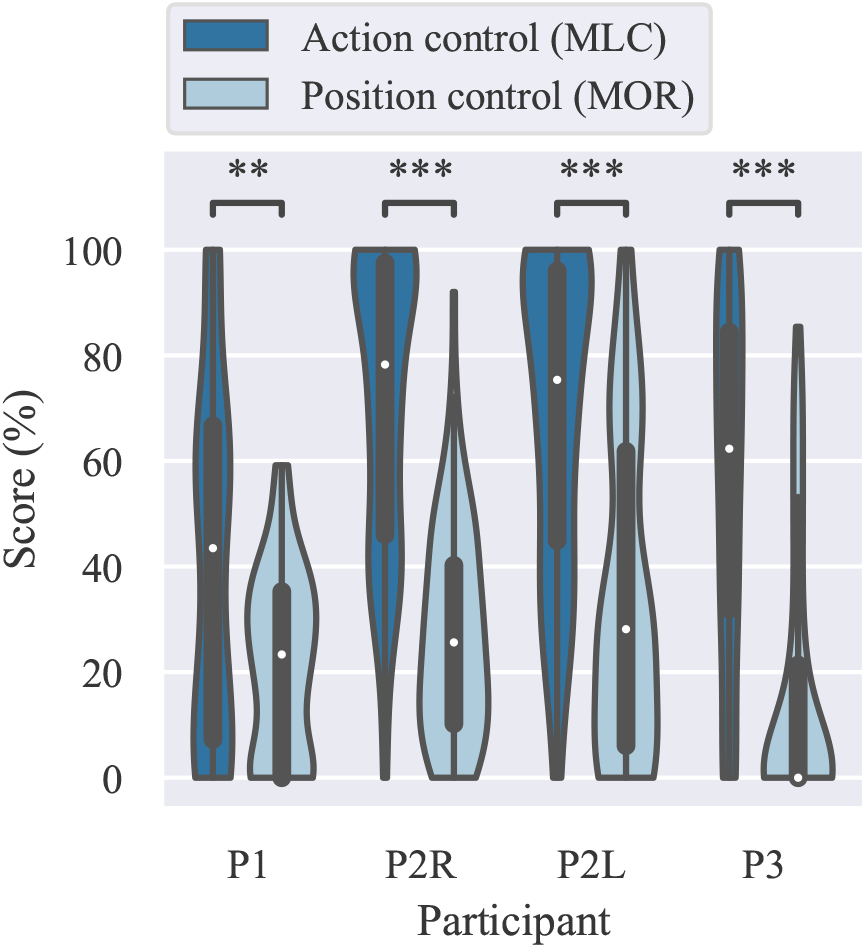
Performance comparison for action (i.e., multi-label classification) and position (i.e., multi-output regression) control. Higher scores indicate better performance. Points, medians; solid boxes, interquartile ranges; whiskers, overall ranges of data; violins, kernel density estimates of underlying data distributions; double asterisk, *p* < 0.01; triple asterisk, *p* < 0.001; MOR, multi-output regression;, MLC, multi-label classification.

A representative trial, performed with each of the two control schemes, is shown in figure 6. The plots show target and user-controlled positions for participant 2R. Position trajectories are juxtaposed for the two control schemes. Target positions are also shown with horizontal black lines. The vertical grey lines indicate the start of the evaluation phase of the trial. The shown example corresponds to the “lateral” grip, for which the thumb rotator is reposed and all digits are fully flexed. It can be observed from this example that action control results in faster target reach, as well as more accurate and stable control. Two video recordings from the same experimental session are provided as supplementary material. They correspond to the last block of trials for each of the two control schemes for the same participant.

**Figure 6:**
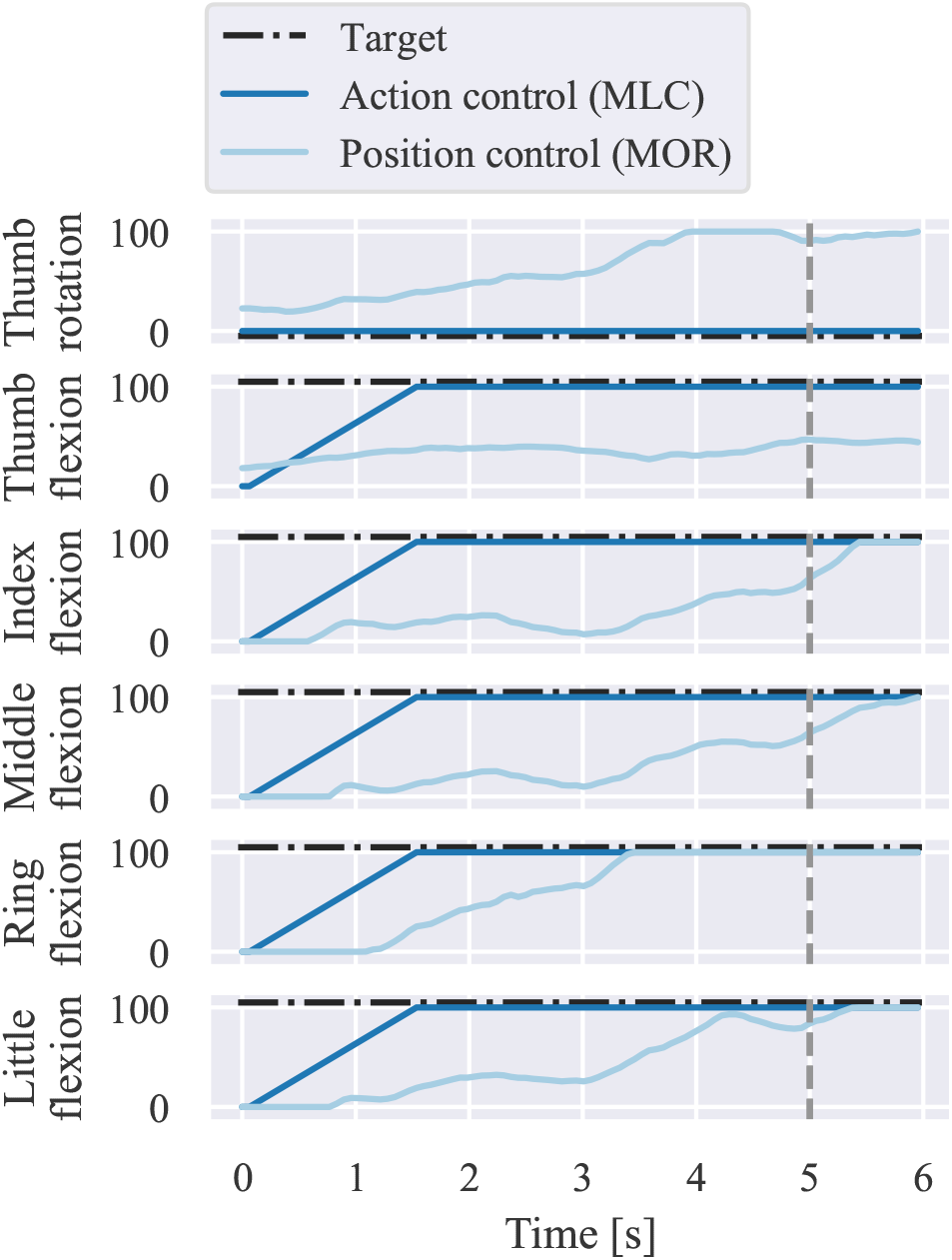
Representative trial. The positions of the six DOFs are juxtaposed for action and position control within the course of one trial. The shown example corresponds to participant 2R and the “lateral grip”, with the following target positions (black dashed lines): full thumb reposition and full flexion of all digits. Vertical grey lines indicate the start of the evaluation phase of the trial. Performance scores for the shown trial were 100% and 46.44% with action and position control, respectively. A small offset has been added to target positions for illustration purposes.

Figure 7 shows DOF-wise, participant-average mean absolute error for the two control schemes. Averages were calculated by combining trials from all blocks and targets. For both controllers, the lowest average error was observed for the little digit DOF. The highest average errors were observed for the middle and thumb digits for action and position control, respectively. For all DOFs, median average errors were lower with action than position control. Statistical tests were not performed due to small sample size (i.e., *n* = 4).

**Figure 7:**
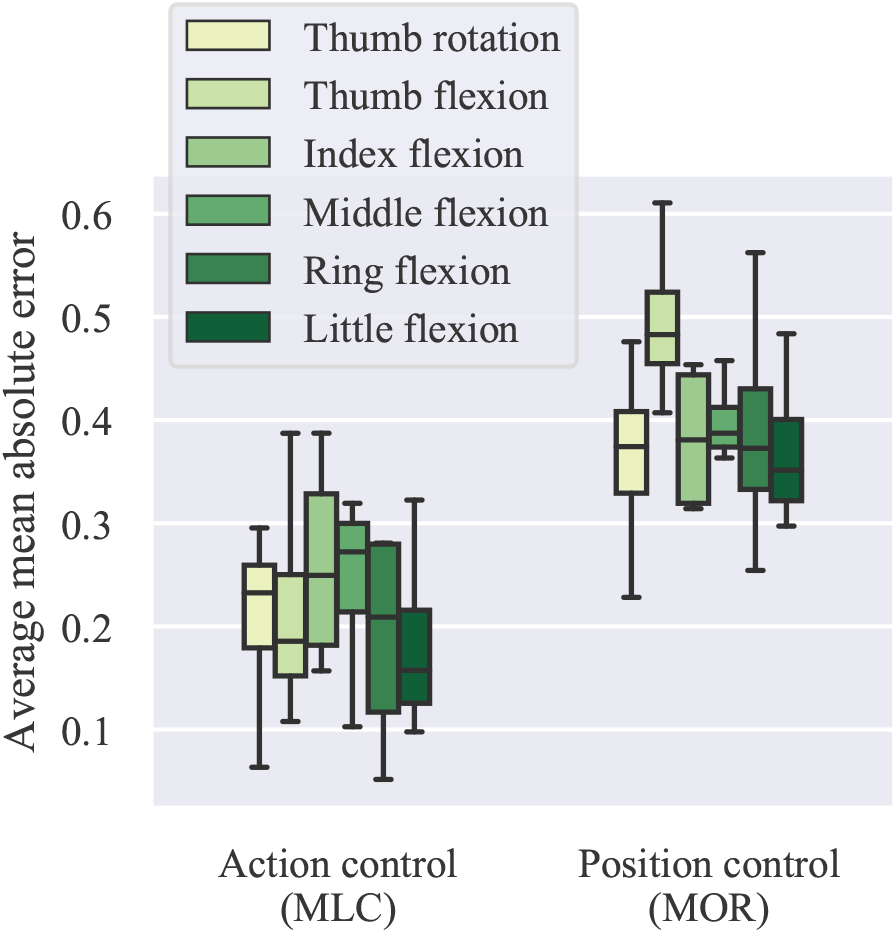
Single-DOF performance comparison. Average mean absolute errors are summarised for action and position control for individual DOFs. Lower scores indicate better performance. Data from all trials are aggregated for each participant. Box plots show distribution of participant-average data (*n* = 4).

We now turn our attention to non-task-related measures. Firstly, we assess muscle contraction intensity levels for the two control schemes. Figure 8(a) shows average raw EMG power across all channels during the whole trial duration. For all participants, action control resulted in significantly lower average raw EMG power (P1, *PMD* = 57.6%, *p* < 10^−10^; P2R, *PMD* = 93.1%, *p* < 10^−15^; P2L, *PMD* = 95.0%, *p* < 10^−15^; P3, *PMD* = 90.2%, *p* < 10^−15^; *PMD* denotes percent median decrease with respect to position control values). In addition, we consider output stability by computing the average output variability (i.e., standard deviation) during the trial evaluation phase. We observed significantly lower output variability with action control for all participants (P1, *PMD* = 70.0%, *p* < 10^−3^; P2R, *PMD* = 92.1%, *p* < 10^−9^; P2L, *PMD* = 86.3%, *p* < 10^−8^; P3, *PMD* = 89.2%, *p* < 10^−15^).

**Figure 8:**
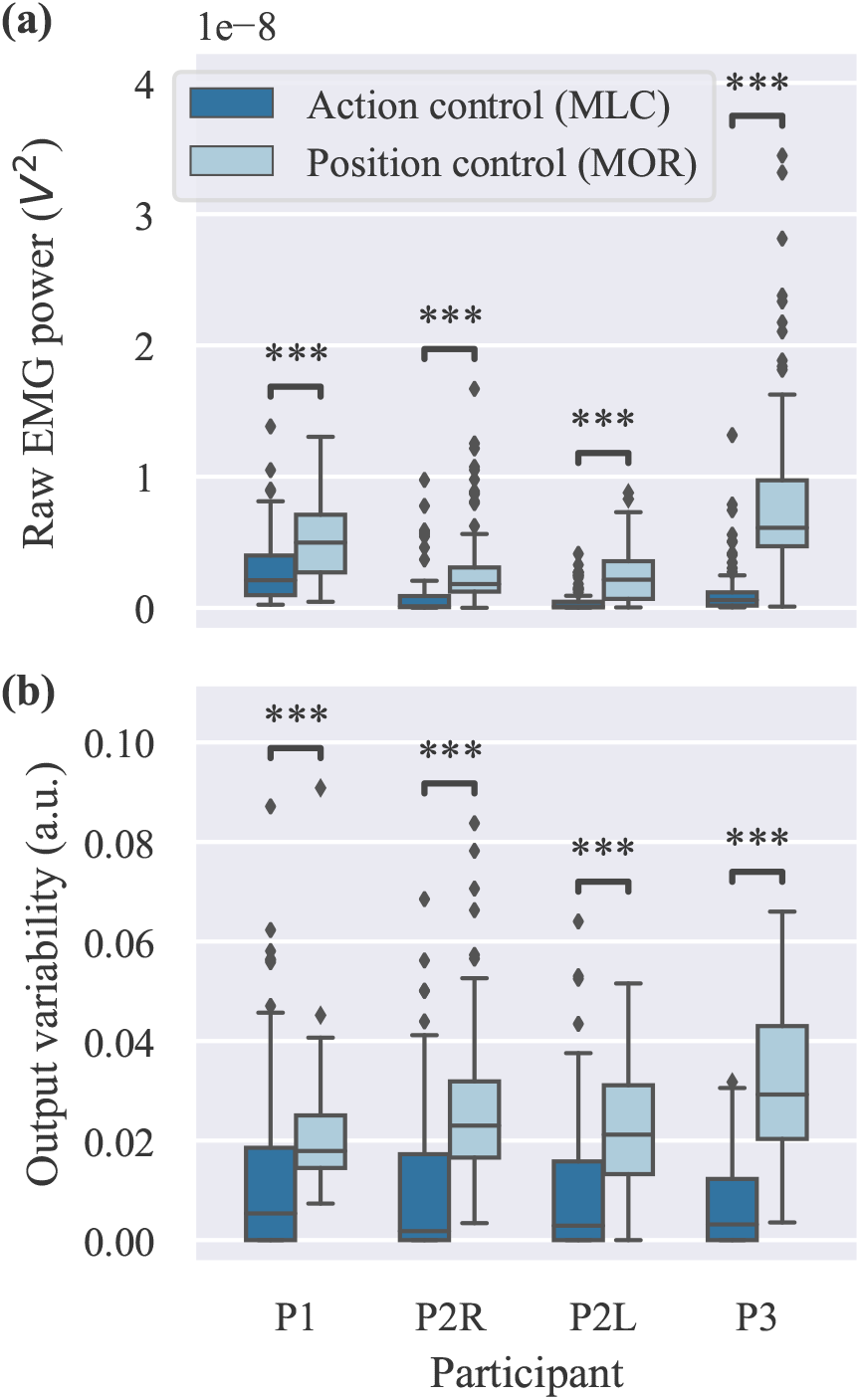
Muscle contraction level and output stability comparison. **(a)** Average raw EMG power for all electrodes during whole trial duration. **(b)** Average output variability during evaluation phase. For each participant, data from all trials are aggregated (*n* = 100 in each box plot). Diamonds indicate outliers.

### 3.3. Post-experimental questionnaire

The outcomes of the post-experimental questionnaire are presented in figure 9. All participants rated action control higher than position control in all three questions. Furthermore, all three participants reported an overall preference for action control.

**Figure 9:**
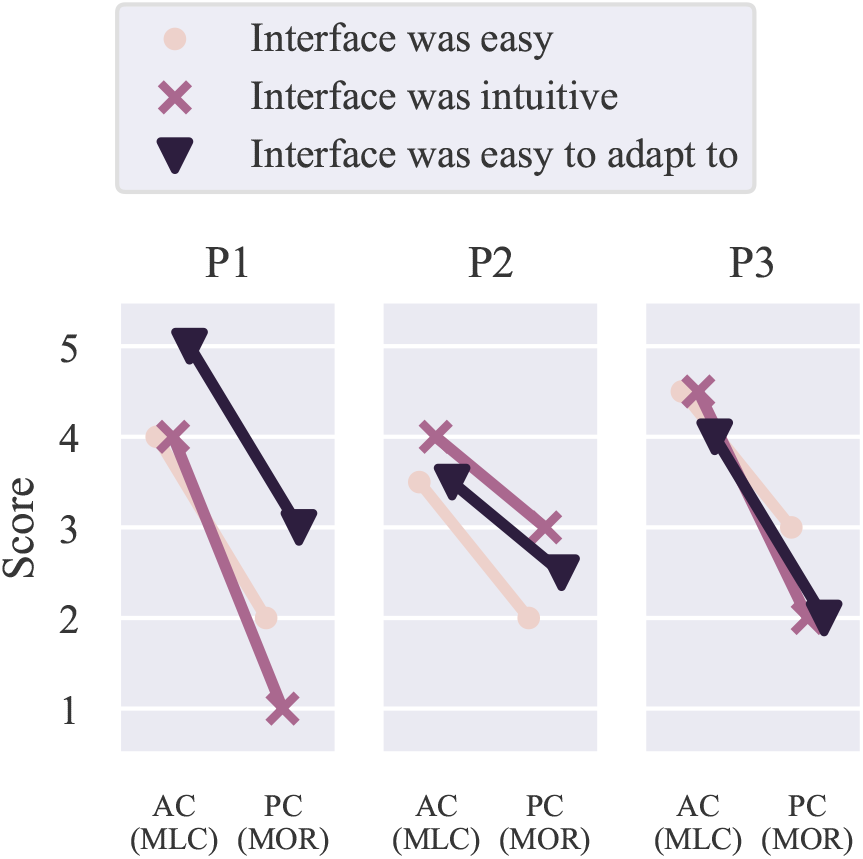
Post-experimental questionnaire. Participants were asked to rate the two control schemes (i.e., action and position control) with three questions. Ratings ranged from 1 (strongly disagree) to 5 (strongly agree). Participant 2 answered the questionnaire twice, once after each session (i.e., right and left sides), and respective scores were averaged. AC, action control; PC, position control.

## 4. Discussion

We have proposed and evaluated *action control*, a novel paradigm for EMG-based prosthesis digit control. The algorithm is based on multi-label, multi-class classification to decode movement intent for each controllable DOF into one of three possible actions: open, close, or stall. We have previously shown that this type of controller can result in comparable performance to direct joint position control in a robotic hand tele-operation task with a data glove [22]. In this study, we have implemented the decoder and control algorithm in real-time using surface EMG measurements and tested it with three transradial amputee participants in a total of four experimental sessions. We have shown that action control can outperform the state-of-the-art approach in myoelectric digit control, which is based on multi-output regression. The observed improvement in performance has been both substantial and systematic across participants. Moreover, it has been systematic across evaluation measures, including both task- and non-task-related metrics. In addition, all participants rated the action control scheme higher than position control in a series of qualitative metrics and reported an overall preference for the former.

In the context of myoelectric control, multi-label (i.e., *simultaneous*) classification has been previously used only to decode wrist and/or whole hand functions [28–32]. For prosthesis digit control, the focus has been on using multi-output regression to decode joint angles (i.e., positions) [9–12,14], velocities [15,16] or fingertip forces [18–20]. Action control can be seen as an extreme, discretised case of velocity control (see figure 2), whereby the velocity can be either zero or take a constant value, which is only parametrised by its sign. Note that, in comparison with classification-based grip control algorithms, generalisation to postures outside the training set is possible with this method. In other words, arbitrary hand configurations are feasible via appropriate sequences of action commands.

A previous study proposed a similar approach to ours [33], however it had three main differences/limitations: 1) it was based on position rather than action control; 2) there were only two classes for each digit, fully open or close, and therefore, intermediate digit positions were not possible; 3) it was limited to offline analysis, and thus, was open-loop. In comparison, the focus of this work was on real-time control with the user in the loop.

It is known and well accepted that interaction with a myoelectric interface that provides the user with some sort of feedback information (e.g., visual), leads to user adaptation over time. When machine learning-based controllers are deployed, this is usually manifested as an increase in task performance [34–36], regardless of the level of intuitiveness of the interface [14]. In fact, there exists a stream of research especially exploiting the high level of plasticity of the human brain to develop myoelectric interfaces that, despite being non-intuitive from a physiological perspective, can be mastered by the user with experience (i.e., *user learning* approach). These are typically not based on machine learning and rather rely on the user’s ability to integrate feedback information to implicitly develop an inverse map of the interface so as to maximise performance in a given task [37–43].

The role of continuous feedback in action control is of particular importance. To achieve the desired digit positions, the user has to estimate the error between the current and target positions and take the appropriate action(s) to minimise it. In other words, the algorithm on its own can be seen as an open-loop control paradigm, in which the user closes the loop by continuously integrating the available feedback information. This requirement might at first seem as a disadvantage in comparison with digit position control. One should keep mind, however, that neither the latter can be considered as entirely biomimetic, mainly due to biomechanical differences between a natural and an artificial limb. As a consequence, the performance of intuitive position control systems also relies heavily on the provision of continuous feedback information [14]. Note that in our experiments the feedback was visual, but it could also take other invasive [44] or non-invasive [45] forms.

In addition to an increase in task performance (figure 5), action control resulted in significantly lower muscle contraction intensity (figure 8(a)) and improved output stability (figure 8(b)). The decrease in EMG amplitude is mainly attributed to the fact that, once the desired posture has been reached, the user can completely relax their muscles. In stark contrast, with position control the user has to hold a muscle contraction in order to retain a desired posture. Lower muscle contraction intensity translates into more effortless control for the user on the one hand, as well as less EMG signal noise on the other, which is known to be signal-dependent [46].

The improvement in control stability is likely due to the discrete nature of the output variable in the case of classification. It is well known that the EMG signal is intrinsically stochastic [47]. In the case of regression, the input noise is propagated through to the output resulting in an unstable controller and unintended prosthesis activations (i.e., jitter effect). To address this issue, it is common to post-process predictions using a low-pass filter (i.e., smoothing) [10, 14, 21]. A large amount of smoothing is typically required to achieve a satisfactory outcome, at the expense of a decrease in the responsiveness of the system. In comparison, a classifier-based controller does not suffer from this limitation, given that the output variables are discrete. The use of a confidence-based rejection strategy further decreases the likelihood of unintended activations [24], hence resulting in more robust and stable digit control. Finally, the increased output stability may lead to further decrease in EMG power, by reducing the amount of needed compensatory contractions.

To train regression-based systems, users are typically instructed to perform bilateral mirrored movements, whilst hand kinematic data are recorded from the contralateral to the EMG side using data gloves [9, 10, 14, 16] or motion tracking systems [11]. This approach has two main limitations: 1) from a clinical perspective, it is highly non-practical; and 2) it can only be applied in the case of unilateral amputation. Indeed, two out of three participants in our study were bilateral amputees. To address this issue, we used computer-generated prompts in combination with linear interpolation to produce regression ground truth labels (section 2.6). However, this approach can introduce label noise, given that it is not guaranteed that participants would precisely follow the prompts with their phantom limb, whilst these are demonstrated on the prosthesis. On the other hand, the use of classification methods avoids this limitation, given that labels are discrete and constant within a trial. The feasibility of collecting noise-free ground truth labels without a requirement for bilateral mirrored training is an additional advantage of the action control approach.

Our study has two limitations. Firstly, participants controlled a computer interface rather than an actual prosthesis. This was due to the lack of position encoders on the prosthesis that we used in our experiments, which were required for benchmarking our proposed algorithm against the regression-based approach. In the future, we will transfer the control interface from the computer display onto the prosthesis and evaluate performance during activities of daily living, such as object picking and manipulation. This will further allow to assess performance robustness under dynamic limb position and prosthesis load conditions. The second limitation was that we used static myoelectric decoders. We plan to extend this work in the future by investigating the potential efficacy of adaptive multi-label classification algorithms.

## 5. Conclusion

We have proposed and evaluated a novel control scheme for myoelectric digit control with transradial amputee participants. Our proposed algorithm, termed *action control*, is based on multi-label, multi-class classification and relies on the provision of continuous visual feedback to the user. We have shown that it can significantly outperform the state-of-the art regression-based approach with respect to several task- and non-task-related measures. In the future, we will further evaluate the performance of the method by transferring the task space from a computer interface onto a real prosthesis.

## Supporting information

Supplementary Movie S1: Action control

Supplementary Movie S2: Position control

## Acknowledgements

The authors are thankful to the three amputee volunteers for their participation. This work is supported by the UK Engineering and Physical Sciences Research Council (EPSRC) under grant EP/R004242/1.

